# CaMKII monomers are sufficient for GluN2B binding, co-condensation, and synaptic potentiation

**DOI:** 10.1101/2025.07.04.663188

**Authors:** Carolyn Nicole Brown, C. Madison Barker, Carley N. Miller, Jason Aoto, Steven J. Coultrap, K. Ulrich Bayer

## Abstract

Cognitive functions require synaptic plasticity, specifically long-term potentiation (LTP). LTP is thought to require CaMKII binding to the NMDA-type glutamate receptor subunit GluN2B, but this poses a major conundrum: Truncated CaMKII monomers (without the hub domain that forms 12meric holoenzymes) fail to bind GluN2B, but still potentiate synapses when made constitutively active. We hypothesized that CaMKII monomer binding to GluN2B has just eluded detection. Instead, even though full-length CaMKII monomers (with hub domain mutations) were found to indeed bind and even co-condensate with GluN2B, truncated monomers were not. Nonetheless, truncated monomers still potentiated synapses, even in neurons with GluN2B mutations that ablate CaMKII binding. However, potentiation occurred only with monomers that were made Ca^2+^-independent by artificial phosphatase-resistant thio-autophosphorylation, not by regular autophosphorylation of T286. These findings support that CaMKII binding to GluN2B is required during physiological LTP induction because it generates the phosphatase-resistant autonomous activity that mediates LTP expression.

## INTRODUCTION

Learning, memory, and other cognitive functions are thought to be mediated by forms of synaptic plasticity, such as long-term potentiation (LTP). It has been established for almost 35 years that LTP requires the neuronal α isozyme of the Ca^2+^/CaM-dependent protein kinase II (CaMKIIα; hereafter referred to as CaMKII)^1–5^. More recently, it was shown that LTP induction specifically requires CaMKII binding to the NMDA-type glutamate receptor subunit GluN2B^6,7^, which also mediates the accumulation of CaMKII at synapses after LTP stimuli^8,9^. LTP expression is then mediated by the Ca^2+^/CaM-independent “autonomous” kinase activity that is generated by CaMKII binding to GluN2B^7,8^. LTP maintenance may still require structural functions of GluN2B-bound CaMKII^10–12^, but kinase activity is no longer required^7^. CaMKII autophosphorylation at T286 (pT286) also generates autonomous CaMKII activity and has long been thought to mediate LTP maintenance^5^, however, pT286 is required only for regulation of GluN2B binding during the initial induction phase^5–7^.

The functional importance of CaMKII binding to GluN2B renewed our interest in exploring some of the remaining conundrums surrounding this binding. Specifically, it is thought that GluN2B binding requires the 12meric CaMKII holoenzyme, as several studies failed to detect binding of immobilized GluN2B to truncated monomeric CaMKII that lacks the C-terminal hub domain that is required for holoenzyme formation^13–15^. However, the N-terminal kinase domain that is still present in the truncated monomers is known to contain a GluN2B binding site that binds to soluble GluN2B peptides^14,16^. Furthermore, several independent sets of classic experiments have shown that constitutively active truncated CaMKII monomers are sufficient to directly cause synaptic potentiation^17–19^. Yet if GluN2B binding is required for LTP induction, and truncated CaMKII monomers do not bind GluN2B, such monomers should not cause LTP.

Here, we set out to test possible explanations for the remaining apparent discrepancies regarding the role of GluN2B binding in LTP. For instance, failure to detect GluN2B binding of CaMKII monomers may not be a binding issue, but a detection issue, as the signal of a bound holoenzyme would be roughly twelve times greater than the signal of a bound monomer. However, we instead detected GluN2B binding for monomeric CaMKII as long as it still contained the hub domain (albeit with mutations to prevent holoenzyme assembly), indicating that efficient binding requires the hub domain. Indeed, such full-length monomers (flMonomers) also moved to excitatory synapses in response to chemical LTP stimuli, whereas truncated monomers (ΔMonomers) did not. Nonetheless, constitutively active ΔMonomers were able to induce synaptic potentiation, even in slices from mice with GluN2B mutations that prevent CaMKII binding. However, this circumvention of GluN2B binding required irreversible constitutive activity by thio-autophosphorylation (thio-pT286) and was not achieved by the reversible pT286 that also occurs naturally during LTP induction. Thus, even though the requirement for CaMKII binding to GluN2B in synaptic potentiation can be circumvented by artificially prolonging autonomous CaMKII activity, physiological LTP requires the autonomous activity of GluN2B-bound CaMKII.

## RESULTS

### Sensitive signal detection in the HEK293T GluN2B binding assay

One potential explanation for the lack of monomer binding to GluN2B in previous studies^13–15^ is simply an issue of detection. Thus, we first examined the detection sensitivity of our established GluN2B binding assay in HEK cells. In this assay, we co-express mEGFP-CaMKII and membrane-targeted (pDisplay) mCherry labelled GluN2B cytoplasmic C-terminus (amino acids 1122-1482 with S1303A mutation; pDisp-mCh-GluN2Bc), induce cellular Ca^2+^ stimuli with ionomycin (10 μM), and then measure the ionomycin-induced change in mEGFP/mCh co-localization as a readout for Ca^2+^/CaM-induced GluN2B binding^8,20,21^. As an mEGFP-tagged CaMKII monomer would have roughly 1/12 the fluorescence signal of an mEGFP-tagged holoenzyme, it may be difficult to detect the signal of monomers co-localized with GluN2B. Thus, we decided to determine if our HEK cell assay is capable of detecting CaMKII binding to GluN2B even when only few of the subunits of the CaMKII holoenzyme are labelled. For this purpose, we used dual color tagged holoenzymes: HEK cells were transfected with different ratios of plasmid coding for CaMKII labelled with either mTurquoise (mTq) or mEYFP in order to express holoenzymes with different ratios of the two distinct fluorophores (Figure 1A). Then, the ratio of expression levels was estimated by the relative fluorescence levels, normalized by the fluorescence generated by a dually labelled mTq-T2A-mEYFP construct that expresses the two fluorescent proteins at a ∼1:1 ratio^22^. We achieved a range of mTq/mEYFP subunit expression ratios from roughly 0.08 to 0.80, where 1.00 is indicative of equal expression of both tagged constructs and where ratios below 1.00 indicate more mEYFP than mTq subunits. Even in cells expressing vastly more mEYFP-CaMKII subunits than mTq-CaMKII subunits, the measured values for the total change in co-localization were virtually identical (Figure 1B). We did not find any correlation between the ratio of mTq/mEYFP subunits and the ratio of the mCh co-localization for each fluorophore (Figure 1C). In fact, for all mTq/mEYFP expression ratios tested, there was robust agreement in the measured change in co-localization with mCh-GluN2Bc for each individually measured fluorophore (Figure 1D). Thus, the number of fluorescent labels per CaMKII holoenzyme does not have an appreciable effect on the detection of GluN2B binding in our HEK cell assay.

**Figure 1.**
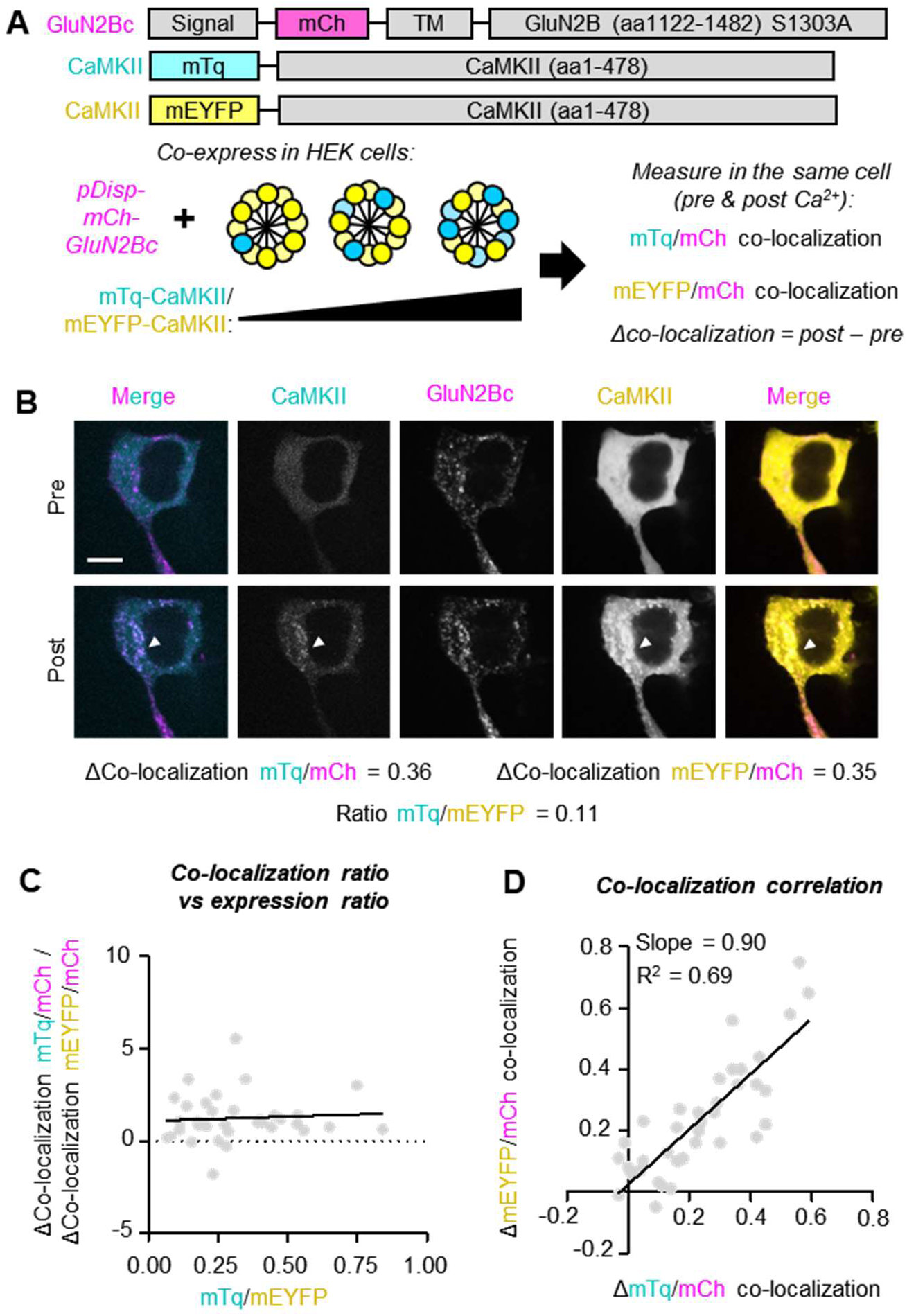
Effects of CaMKII signal on reported GluN2B co-localization. **(A)** Schematic of experiment shown in B) and C). mTq-CaMKII and mEYFP-CaMKII were co-expressed using different [DNA] ratios to achieve formation of holoenzymes with a mixture of mTq and mEYFP subunit tags. Each fluorophore was quantified for total fluorescence intensity and actual ratios were estimated based on fluorescence intensity of a control plasmid that expresses both fluorophores at a 1:1 ratio. Both mTq-CaMKII and mEYFP-CaMKII were separately quantified for their co-localization with pDisp-mCh-GluN2Bc. **(B)** Representative images of a cell expressing holoenzymes with a majority of mEYFP subunits. The measured total change in ionomycin-induced GluN2B binding (ΔCo-localization; post – pre) is very similar regardless of whether it is quantified for co-localization with the mTq or the mEYFP subunits. **(C)** ΔCo-localization ratio (ΔCo-localization mTq/mCh / ΔCo-localization mEYFP/mCh) versus fluorescence ratio (mTq/mEYFP). A ΔCo-localization ratio above 1 indicates higher measured binding for the mTq-tagged subunits while a ratio below 1 indicates higher measured binding for the mEYFP-tagged subunits. The fluorescence ratio is plotted where 1.0 would indicate a 1:1 measured expression ratio and <1.0 would indicate more mEYFP-CaMKII than mTq-CaMKII expression. **(D)** Shown here is the ΔCo-localization for mEYFP/mCh versus the ΔCo-localization for mTq/mCh.

### Validation of two distinct monomeric CaMKII constructs

With successful detection of CaMKII holoenzyme binding to GluN2B even with sparsely labelled holoenzymes, we next revisited detection of binding for CaMKII monomers. In addition to the previously published truncated monomers^13–15^, we developed a full-length monomer that still contains the hub domain, but with point mutations that prevent holoenzyme assembly. The truncated monomeric construct was made by placing a stop codon before the CaMKII hub (317stop; ΔMonomer) (Figure 2A), similar as in previous studies. The second construct received point mutations in the horizontal and vertical interfaces of the CaMKII hub to prevent formation of holoenzymes (F394A H468E; flMonomer) (Figure 2A). We validated the monomeric nature of both mEGFP-tagged constructs by three independent assays. First, we used *in vitro* fluorescence correlation spectroscopy (FCS) to measure the molecular brightness (Figure 2B) and diffusion times (Figure 2C), as we have described previously^15,21,23^. Both the flMonomer and the ΔMonomer had significantly decreased molecular brightness and diffusion time compared to WT CaMKII (“Holoenzyme”) but were not different from each other or from a non-conjugated mEGFP.

**Figure 2.**
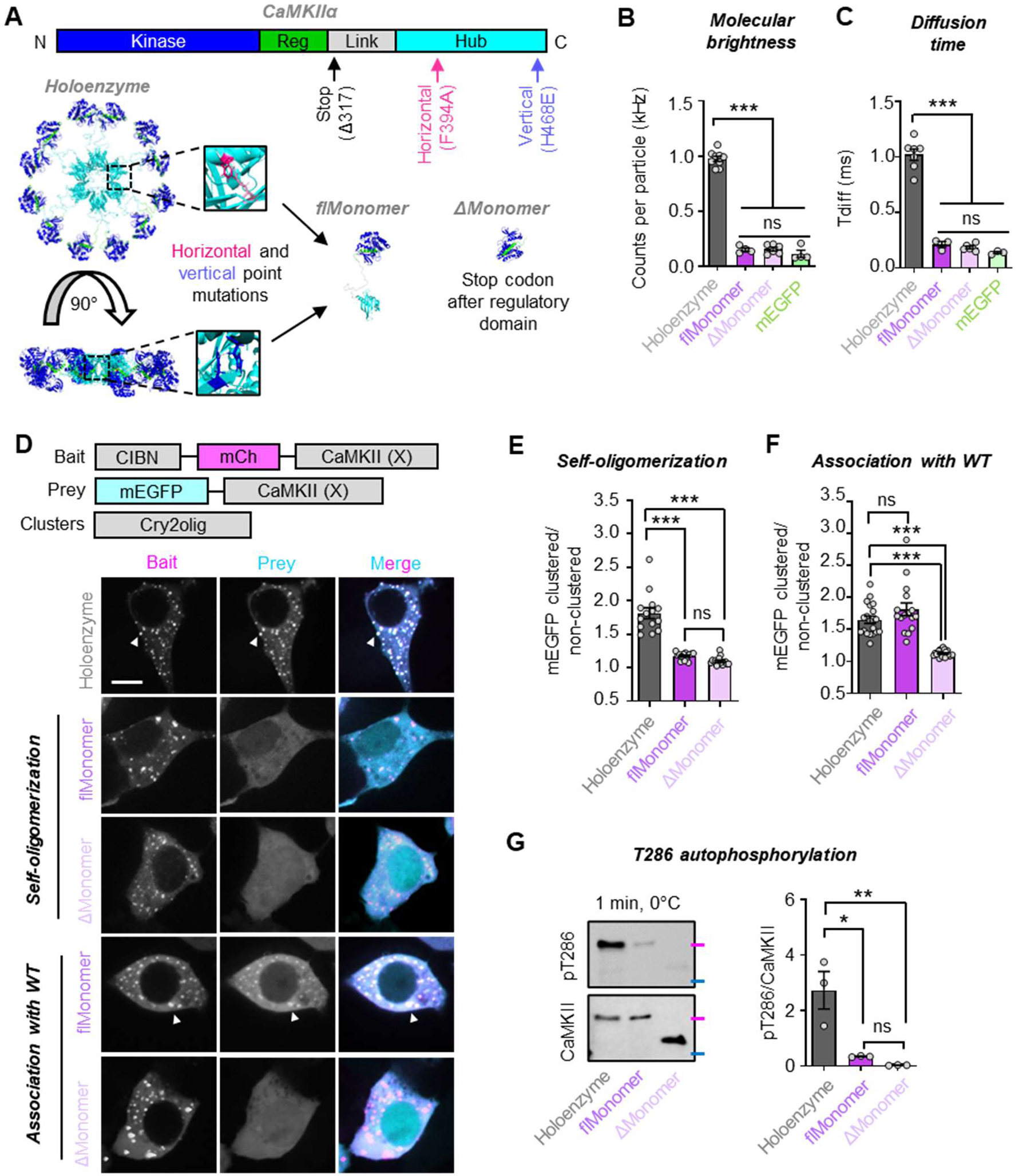
Generation and validation of CaMKII full-length monomer and truncated monomer. **(A)** Primary sequence and structure of WT CaMKII holoenzyme. Point mutations were strategically made in the hub horizontal and vertical interfaces to generate a full-length, monomeric CaMKII (flMonomer; F394A H468E). The truncated monomer (ΔMonomer) was generated by placing a stop codon after the regulatory domain (317stop). Fluorescence correlation spectroscopy was used to measure **(B)** the molecular brightness and **(C)** the diffusion time of the two monomeric constructs versus the holoenzyme and the mEGFP tag itself. ***P<0.001; ns, not significant by one-way ANOVA. **(D)** Constructs used to assay self-oligomerization and WT association of monomeric mutants. **(E)** Here, monomeric mutants were expressed on both Prey (mEGFP) and Bait (CIBN-mCh) backgrounds to assay for self-oligomerization or **(F)** monomeric mutants (X) were expressed as Prey (mEGFP) with WT holoenzymes as Bait (CIBN-mCh) to assay for association with WT CaMKII. ***P<0.001, ns, not significant by one-way ANOVA. Scale bar = 10 μm. **(G)** *In vitro* kinase reactions (50 mM PIPES pH 7.2, 2 mM Ca^2+^, 10 mM Mg^2+^, 1 mM ATP, 1 μM CaM) were performed on ice (0°C) for 1 minute with 5 nM subunit concentration for each mEGFP-tagged construct isolated from HEK293T cells. Molecular weight markers are color coded based on the BioRad Kaleidoscope Standard: magenta = 75 kDa, blue = 50 kDa. *P<0.05; **P<0.01; ns, not significant by one-way ANOVA.

To further confirm the monomeric nature of these constructs also within cells, we utilized an established light-induced co-clustering (LINC) assay^15,24,25^. First, we expressed each CaMKII construct on both the bait (CIBN-mCh-CaMKII) and prey (mEGFP-CaMKII) backgrounds to probe for self-oligomerization (Figure 2D,E). As expected, the holoenzyme displayed robust co-clustering upon light stimulation, indicative of its self-oligomerization. In contrast, no co-clustering was detected for mEGFP-flMonomer when co-expressed with CIBN-mCh-flMonomer, nor for mEGFP-ΔMonomer when co-expressed with CIBN-mCh-ΔMonomer. This indicates that both the flMonomer and ΔMonomer are indeed monomeric when expressed at high concentrations within cells. We additionally tested whether either monomeric construct associates with WT CaMKII (Figure 2D,F). To this end, we expressed WT CIBN-mCh-CaMKII (WT) with mEGFP-tagged monomeric constructs. Here, we still found no evidence of mEGFP-ΔMonomer association with WT CIBN-mCh-Holoenzyme. In contrast, we detected association of mEGFP-flMonomer with WT CIBN-mCh-Holoenzyme (Figure 2F). Thus, while we confirmed that the full-length monomeric construct is indeed monomeric when expressed alone, it retains its ability to interact with WT CaMKII. This needs to be taken into consideration when performing experiments in neurons, where a significant amount of endogenous WT CaMKII is present.

Finally, we confirmed the monomeric nature of our CaMKII mutants by an *in vitro* pT286 assay. While pT286 has been demonstrated as both an intra- and inter-holoenzyme reaction, it does not occur as an inter-holoenzyme reaction at low concentrations of CaMKII^26–28^. Thus, at low concentrations, pT286 should be significantly impaired for any monomeric CaMKII. Indeed, both mEGFP-flMonomer and mEGFP-ΔMonomer were significantly impaired for pT286 compared to mEGFP-Holoenzyme, but did not significantly differ from each other (Figure 2G). Thus, three independent lines of evidence confirmed that both the flMonomer and the ΔMonomer are indeed monomeric.

### GluN2B binding detection for the CaMKII monomer that includes the hub domain

Next, we compared GluN2B binding of CaMKII holoenzymes and our two different monomeric CaMKII mutants. To this end, we used the same HEK cell binding assay as in Figure 1. Again, the S1303A GluN2B mutation is used in order to eliminate this CaMKII phosphorylation site that negatively regulates CaMKII binding; in turn, this eliminates possible confounds in case of differential phosphorylation by the three distinct CaMKII forms. As expected, the wildtype CaMKII holoenzyme showed a robust increase in GluN2B binding in response to the ionomycin-induced Ca^2+^ stimulus. By contrast, the ΔMonomer showed no GluN2B binding (Figure 3B,C), consistent with previous studies with truncated monomers^13–15^. However, for the flMonomer, robust Ca^2+^-dependent binding was detected (Figure 3B), even though the extent was less compared to holoenzymes (Figure 3C). This successful detection of GluN2B binding for the flMonomer further supports our earlier conclusion that there is no issue of detecting GluN2B-bound CaMKII monomers in principle (see Figure 1). Most importantly, the GluN2B binding of the full-length flMonomer but not the truncated ΔMonomer suggested that the CaMKII hub domain is a positive regulator of GluN2B binding.

**Figure 3.**
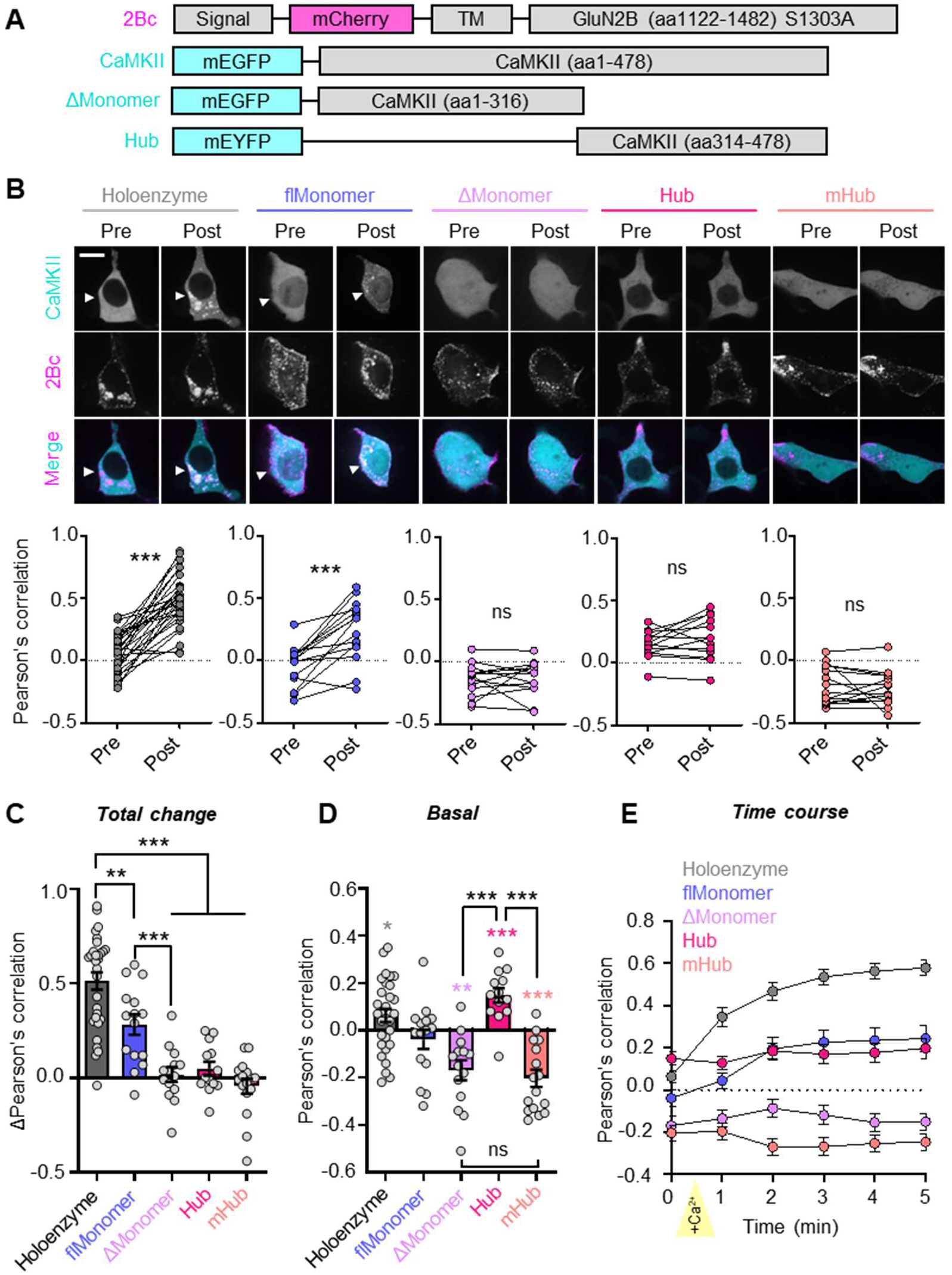
CaMKII hub positively regulates Ca^2+^-dependent GluN2B binding in HEK293T cells. **(A)** Constructs used in this experiment. flMonomer = F394AH468E, ΔMonomer = 317stop, mHub = Hub + F394AH468E. **(B)** Representative images and paired quantification of CaMKII-GluN2B co-localization by Pearson’s correlation pre- and 5 minutes post-stimulation with ionomycin (10 μM). ***P<0.001; ns, not significant by paired t-test. Scale bar = 15 μm. **(C)** Total change in co-localization (post – pre) upon ionomycin stimulation. **P<0.01, ***P<0.001, one-way ANOVA. **(D)** CaMKII-GluN2B co-localization pre-ionomycin stimulation. Colored asterisks: *P<0.05, **P<0.01, ***P<0.001 vs 0, one sample t-test. Black asterisks: ***P<0.001, one-way ANOVA. **(E)** Time course of CaMKII-GluN2B co-localization. Yellow arrow indicates start of ionomycin treatment (“+Ca^2+^”).

To examine a role for the hub domain as a positive regulator of GluN2B binding, we generated constructs that express only the oligomerization-competent hub domain of wild-type CaMKII (“Hub”) or a monomeric mutant (“mHub”; Hub + F394A HA468E). For the hub domain alone, we found no indication for Ca^2+^-induced GluN2B binding, neither for the 12meric Hub nor for its monomeric mHub counterpart (Figure 3B,C). However, before the Ca^2+^ stimulation, we found basal co-localization with GluN2B for the 12meric Hub and basal exclusion from GluN2B for the monomeric mHub (Figure 3D). We also observed basal co-localization for the full-length flMonomer and basal exclusion for the ΔMonomer with GluN2B. Additionally, for all three monomeric constructs (flMonomer, ΔMonomer, mHub), we observed partial nuclear localization (Figure 3B), as expected based on less stringent nuclear exclusion of smaller proteins and further confirming their monomeric nature. Additionally, we determined that there was no effect of the fluorescent protein tags used (mEGFP, mEYFP) on GluN2B binding in our HEK cell assay and neither fluorophore showed any GluN2B co-localization on its own (Supplemental Figure S1A,B). Similar to the mEYFP-Hub (Figure 3), we found that the mEGFP-Hub showed mild, but significant basal co-localization with GluN2B and lacks any Ca^2+^-dependent increase in co-localization (Supplemental Figure S1E-G).

Overall, the results from our HEK cell binding assay suggest a Ca^2+^-independent interaction of the CaMKII hub with GluN2B and showed that inclusion of the hub domain allows efficient Ca^2+^-induced GluN2B binding also for monomeric CaMKII. At the same time, they confirm previous studies that did not detect GluN2B binding of truncated monomeric CaMKII that lacked the hub domain^13–15^.

### The CaMKII monomer with hub domain can co-condensate with GluN2B

In addition to traditional high-affinity binding, CaMKII and the cytoplasmic GluN2B C-tail have been recently shown to undergo liquid-liquid phase separation (LLPS), a form of biomolecular co-condensation that is mediated by multiple weak binding interactions and that is proposed to contribute to structural organization of synapses^29–33^. Similar to traditional binding, CaMKII co-condensation with GluN2B is induced by Ca^2+^/CaM-stimuli and was not detected for truncated CaMKII monomers^29^. For co-condensation, it seemed likely that the multivalent nature of CaMKII holoenzymes is required for the multiple interactions that are required for condensation^29,34^. However, as our findings suggested multiple interactions with GluN2Bc even for a full-length CaMKII monomer (i.e., via kinase domain and via hub domain), we decided to compare our two distinct monomers also for co-condensation with GluN2B. Although CaMKII co-condensation with GluN2B has been mainly studied in biochemical assays with purified protein, we have recently developed a co-condensation assay within HEK cells^7,35^. The assay is very similar to our traditional binding assay in HEK cells (in Figures 1 and 3), but instead of a membrane-targeted mCh-GluN2Bc, we use a soluble version of mScarlet-GluN2Bc (Figure 4A) that is distributed throughout the cytoplasm, similar to mEGFP-CaMKII^7^. However, when a Ca^2+^ signal is induced by ionomycin (10 μM), co-condensation of mEGFP-CaMKII with the soluble mScarlet-GluN2Bc can be measured by formation of mEGFP puncta that also contain mScarlet (Figure 4B). Once these co-condensates are induced by Ca^2+^/CaM-stimulation, they persist even when Ca^2+^ is removed by addition of EGTA (Figure 4B). Consistent with prior biochemical assays, our truncated ΔMonomer failed to form co-condensates in this HEK cell assay (Figure 4B, C). By contrast, our flMonomer that contains the hub domain formed co-condensates with GluN2B upon stimulation with ionomycin; once formed, these co-condensates were maintained even after removal of Ca^2+^ with EGTA (Figure 4B). Thus, in contrast to the ΔMonomer, the flMonomer showed overall similar co-condensation with GluN2B as the CaMKII holoenzyme (Figure 4C). However, compared to the CaMKII holoenzyme, the flMonomer already showed some basal co-condensation, even though this was significantly further enhanced by the stimuli (Figure 4D). Together, these results further support that there are indeed multiple binding sites on each CaMKII subunit and on GluN2Bc; unexpectedly, they demonstrate that even monomeric CaMKII can co-condensate with GluN2Bc as long as it is a full-length monomer that also contains the hub domain.

**Figure 4.**
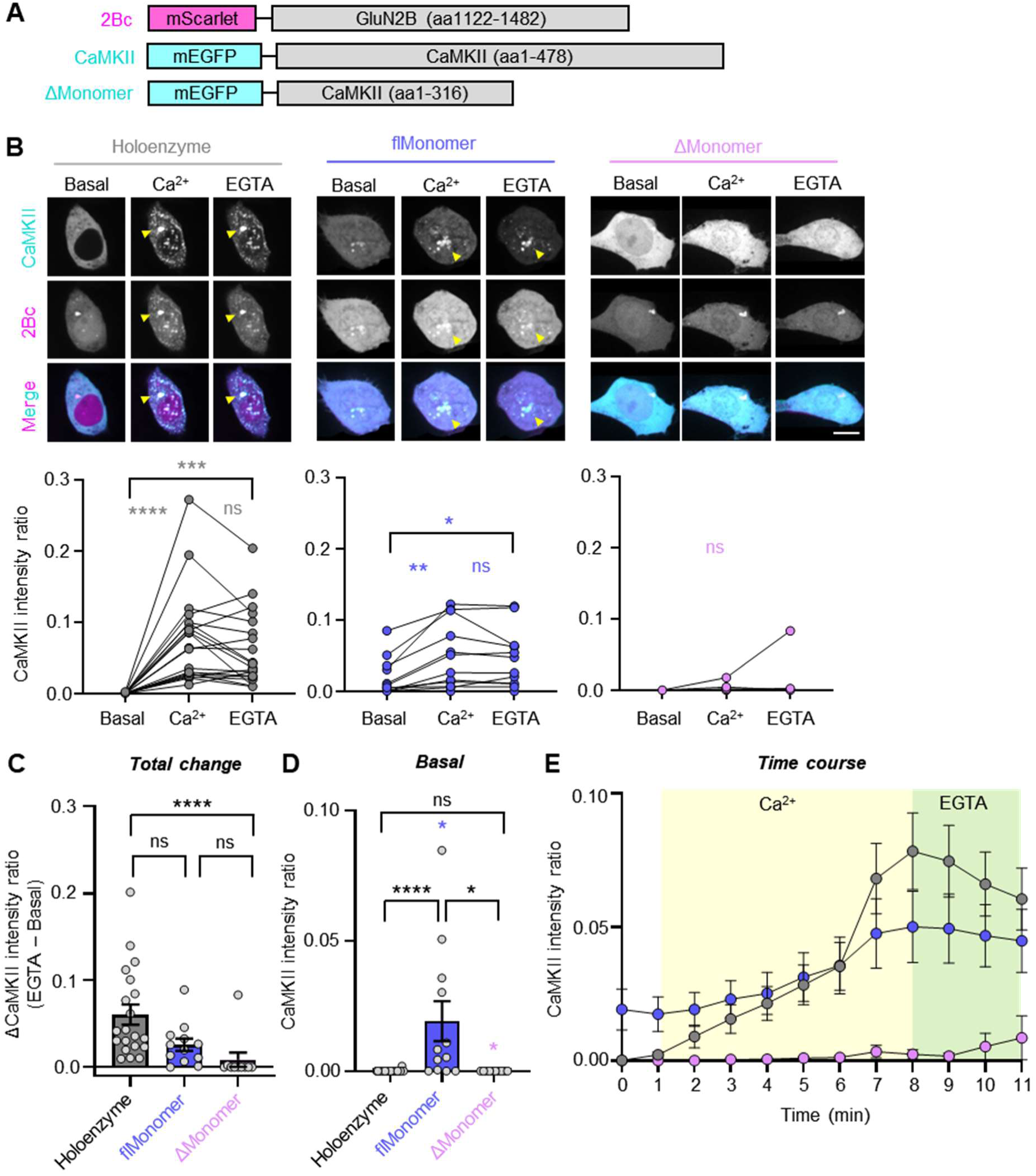
CaMKII full-length monomers are sufficient for co-condensation with GluN2B in HEK293T cells. **(A)** Constructs used in this experiment. flMonomer = F394AH468E, ΔMonomer = 317stop. **(B)** Representative images and paired quantification of mEGFP-CaMKII and mScarlet-GluN2Bc basally, after stimulation with ionomycin (10 μM), and after treatment with EGTA (2.5 mM). Scale bar = 10 μm. *P<0.05; **P<0.01; ***P<0.001; ****P<0.0001; ns, not significant by RM one-way ANOVA. **(C)** Basal CaMKII intensity ratio. *P<0.05; ****P<0.0001; ns, not significant by one-way ANOVA. **(D)** Total change in CaMKII intensity ratio (EGTA – basal). ****P<0.0001; ns, not significant by one-way ANOVA. **(E)** Time course quantification of co-condensate formation between CaMKII and GluN2B.

### The CaMKII hub domain enhances GluN2B binding *in vitro*

Next, we used purified GST-GluN2Bc that was immobilized in microtiter plate wells to compare binding of the flMonomer and ΔMonomer in our established biochemical *in vitro* binding assay^6–8^. Binding was induced with Ca^2+^/CaM (2 mM/1 μM) in the presence nucleotide (100 μM ADP) and then measured by western blot after eluting the bound protein from the plates (Figure 5). We observed a significantly greater GluN2B binding signal for the flMonomer compared to the ΔMonomer (Figure 5A,B). The residual binding signal detected for the ΔMonomer was mostly background, as there was no significant difference in presence of Ca^2+^/CaM compared to EGTA (Figure 5C).

**Figure 5.**
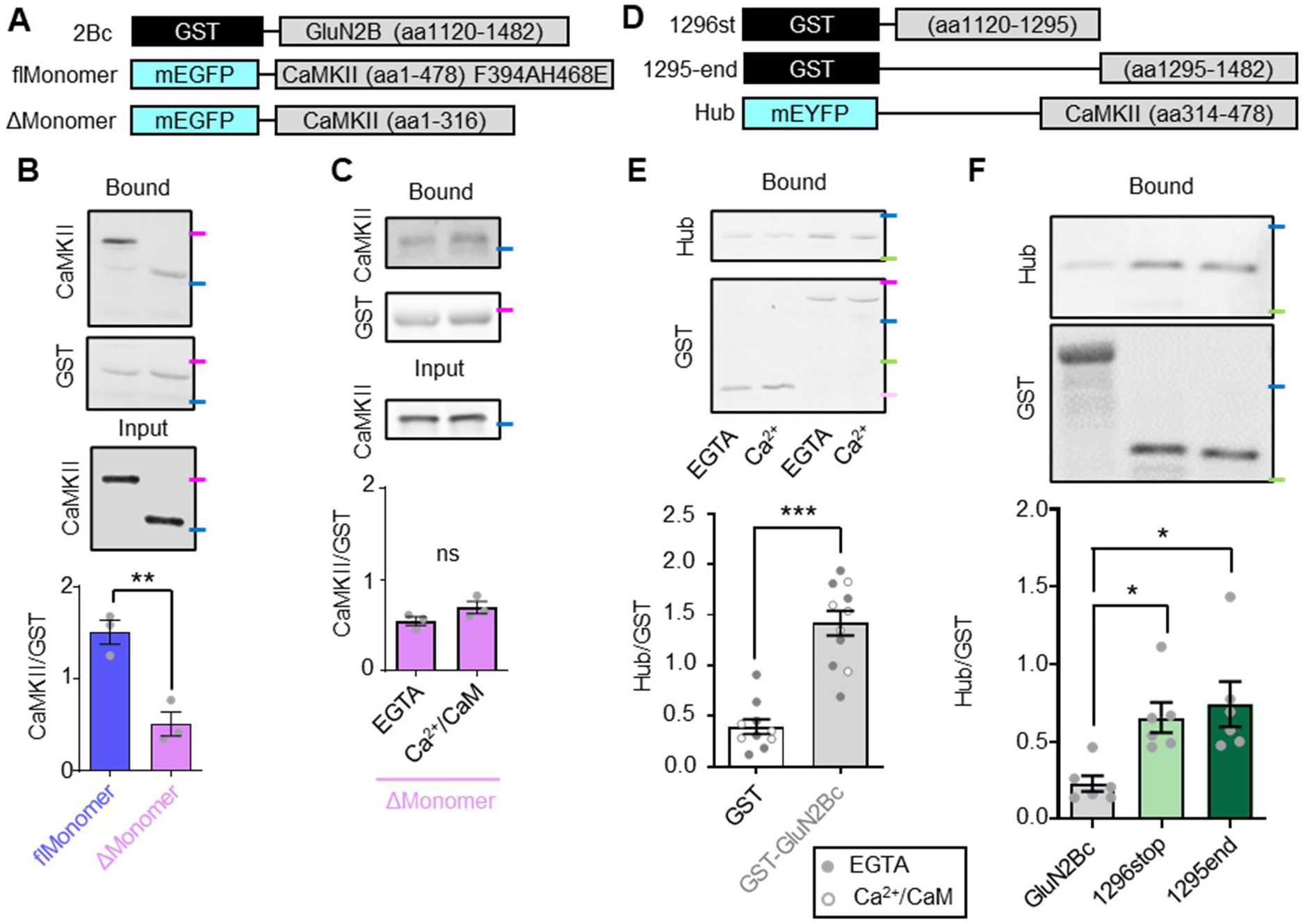
Binding of the CaMKII 12meric hub domain to GluN2B *in vitro*. **(A)** Constructs used to test whether the hub domain enhances GluN2B binding *in vitro*. flMonomer = F394AH468E, ΔMonomer = 317stop. **(B)** *In vitro* GluN2B binding of flMonomer versus ΔMonomer in the presence of Ca^2+^/CaM (2 mM/1 μM) and ADP (100 μM). **P<0.01 by unpaired t-test. **(C)** *In vitro* GluN2B binding of ΔMonomer in the presence of EGTA (1 mM) versus Ca^2+^/CaM (2 mM/1 μM). ns, not significant by unpaired t-test. **(D)** Constructs used to test for direct binding of the CaMKII 12meric hub to GluN2B *in vitro*. The full-length GST-GluN2Bc construct is the same as shown in A). **(E)** Binding of the hub domain to GST alone vs GST-GluN2Bc. Open circles represent experiment in the presence of Ca^2+^/CaM (2 mM/1 μM) and filled circles represent experiment in the presence of EGTA (1 mM). ***P<0.001 by unpaired t-test. **(F)** Binding of the hub domain to GST-GluN2Bc, GST-GluN2Bc 1296stop, or GST-GluN2Bc 1295-end in the presence of EGTA (1 mM). *P<0.05 by one-way ANOVA. Molecular weight markers are color coded based on the BioRad Kaleidoscope Standard: magenta = 75 kDa, blue = 50 kDa, green = 37 kDa, light pink = 25 kDa.

While we determined that the CaMKII hub alone cannot mediate Ca^2+^-induced GluN2B binding in HEK cells, the enhanced basal co-localization of the hub with GluN2Bc in HEK293T cells (Figure 3D) suggested that there could be direct binding of the hub even in the absence of Ca^2+^. Indeed, in our biochemical *in vitro* binding assay, the CaMKII hub bound to GST-GluN2Bc significantly more than to GST protein alone, indicating direct binding (Figure 5D, E). Consistent with the observation in HEK cells, the *in vitro* binding was Ca^2+^-independent, as it was detected also in the presence of EGTA.

Next, we sought to identify the region of GluN2Bc bound by the hub. Thus, we generated two fragments of GST-GluN2Bc around the kinase binding site: GST-GluN2Bc 1296stop and GST-GluN2Bc 1295-end (Figure 5D). To our surprise, we found that the hub bound equally to both fragments and that this binding was enhanced compared with binding to the full-length GST-GluN2Bc construct (Figure 5F). This result suggested that the hub binding site is partially occluded in GluN2Bc and that there are multiple hub domain binding sites on GluN2Bc. Together, the findings of our *in vitro* binding assay further support that the CaMKII interaction with GluN2B is mediated by at least two binding sites on each CaMKII subunit (one on the kinase and one on the hub domain) and on each GluN2B C-tail, consistent with the successful co-condensation even of the monomeric mCh-GluN2Bc with the CaMKII flMonomer. Thereby, our *in vitro* assays also confirm the necessity of the hub domain for both GluN2B binding and co-condensation.

### A neuroprotective γ-hydroxybutyrate analog does not affect hub binding to GluN2B

The γ-hydroxybutyrate analog 3-hydroxycyclopent-1-enecarboxylic acid (HOCPCA) binds to the CaMKIIα hub domain and protects neurons from excitotoxic or ischemic cell death^36,37^. However, the neuroprotective mechanism remains unclear, as HOCPCA does not inhibit enzymatic CaMKII activity^35,36^. A proposed alternative mechanism was inhibition of GluN2B binding^35,36,38^, which would indeed be neuroprotective^39^. However, we have recently shown that HOCPCA does not prevent GluN2B binding, at least for the CaMKII holoenzyme^35^. Here, we found that HOCPCA does also not prevent the basal, Ca^2+^-independent GluN2B binding of the isolated hub domain assembly (Supplemental Figure S2). Thus, the neuroprotective effect of HOCPCA is not mediated by a direct effect on CaMKII binding to GluN2B.

### The CaMKII hub domain enables cLTP-induced synaptic targeting of monomers

After assessing binding in HEK cells and *in vitro*, we next compared the synaptic accumulation of monomeric and WT CaMKII in hippocampal neurons after chemical LTP stimuli (cLTP; 100 μM glutamate, 10 μM glycine for 30 sec, followed by washout), a process that is mediated by Ca^2+^/CaM-induced GluN2B binding^8,9^ (Figure 6). Endogenous CaMKIIα was knocked down by shRNA, as we have previously described^21,40^. This knock down reduces overall level of CaMKII overexpression and, more importantly, minimizes the interaction of the flMonomer with WT CaMKII that was indicated by our LINC assay (see Figure 2F). All CaMKII constructs tested showed basal synaptic enrichment, with spine to shaft ratios greater than one (Figure 6B,C). Although the enrichment appeared greater for the CaMKII holoenzyme, this was statistically significant only in comparison to the two hub domain constructs without kinase domains (Figure 6C). Then, cLTP stimuli caused significant further synaptic accumulation of the CaMKII flMonomer that was indistinguishable from WT CaMKII (Figure 6B,D). By contrast, the ΔMonomer and the two hub domain constructs without kinase domain (Hub, mHub) failed to show any significant synaptic accumulation in response to the cLTP stimuli (Figure 6B,D), as also evident in the quantification of enrichment during the 5 min time course of the live imaging (Figure 6E). Overall, our results demonstrate that the CaMKII hub domain promotes GluN2B binding *in vitro*, in HEK cells, and in neurons.

**Figure 6.**
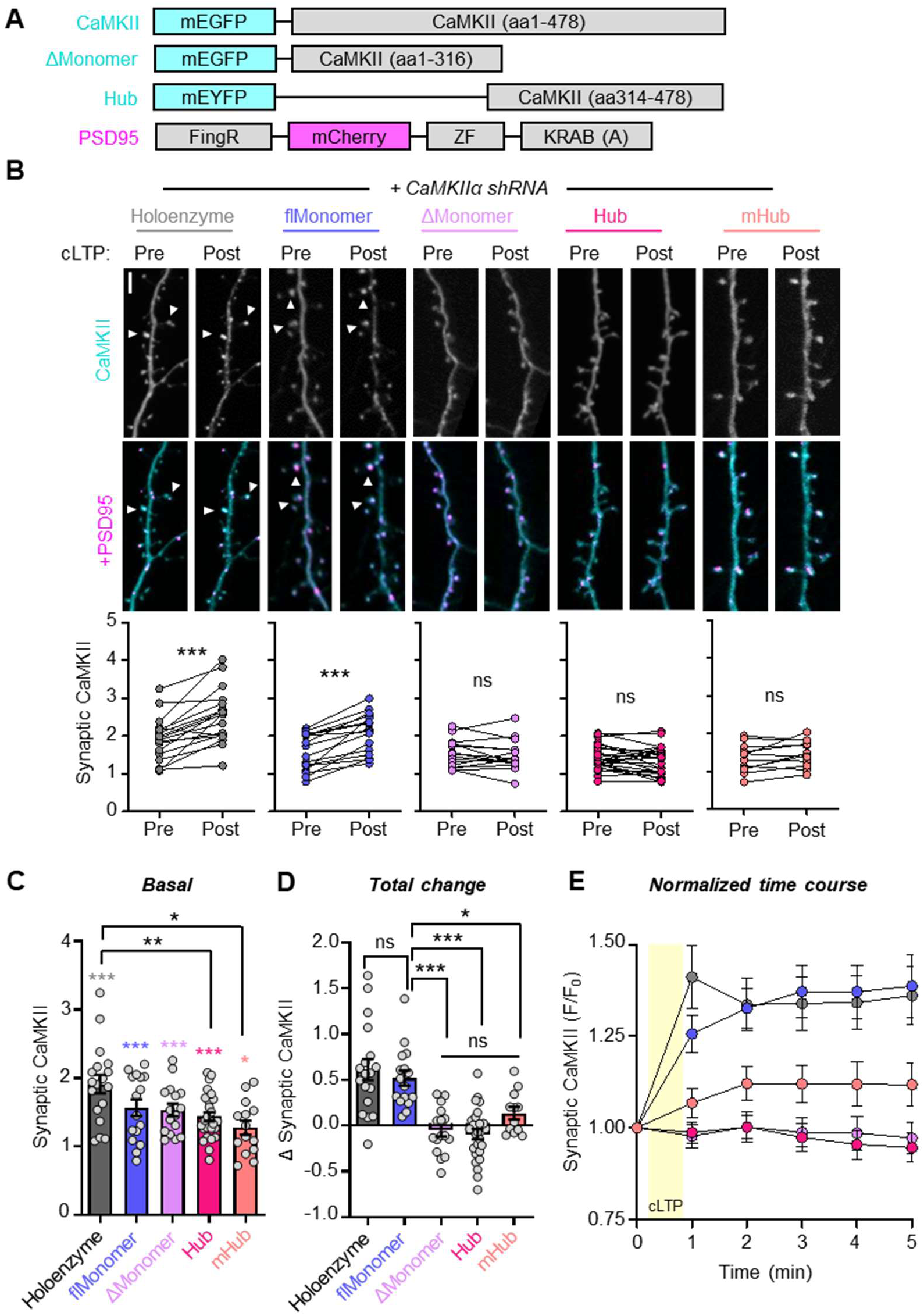
CaMKII hub positively regulates cLTP-induced synaptic localization. **(A)** Constructs used in this experiment. Endogenous CaMKIIα was knocked down with CaMKIIα shRNA in every condition. flMonomer = F394AH468E, ΔMonomer = 317stop, mHub = Hub + F394AH468E. **(B)** Representative images and paired quantification of CaMKII localization pre- and 5 min post-cLTP (100 μM glutamate, 10 μM glycine, 30 sec). ***P<0.001; ns, not significant by paired t-test. Scale bar = 5 μm. **(C)** Basal synaptic CaMKII (pre). Colored symbols: *P<0.05; ***P<0.001 vs 1 with one sample t-test. Black symbols: *P<0.05; **P<0.01 by one-way ANOVA. **(D)** Total change (post – pre synaptic CaMKII). *P<0.05; ***P<0.001; ns, not significant by one-way ANOVA. **(E)** Time course of CaMKII synaptic localization normalized to baseline synaptic localization value (F/F_0_). Yellow area indicates duration of cLTP treatment.

### A constitutively active CaMKII ΔMonomer causes synaptic potentiation without GluN2B binding

The results from this study show that even monomeric CaMKII can bind to GluN2B if it contains a hub domain, but also confirm that truncated monomers that lack the hub domain do not show detectable binding. But then, why can constitutively active truncated monomers mediate potentiation^17–19^ if LTP requires CaMKII binding to GluN2B^6,7,9,10,12^? In order to address this question, we decided to first repeat one of the classic experiments in our hands, specifically by using a purified truncated monomer [6xHis-SUMO (6HS)-CaMKII 317stop] that was made constitutively active by thio-autophosphorylation (thio-pT286; Figure 7A). Indeed, as in the original publication^18^, we found that the thio-pT286 ΔMonomer caused significant potentiation of CA3-CA1 synaptic transmission when introduced into CA1 cells by patch pipette (Figure 7B). After heat-inactivation, the same thio-pT286 ΔMonomer protein preparation did not cause any potentiation, as expected. Next, we tested if the potentiation may be due to residual GluN2B binding of the truncated monomer that eluded our detection: we repeated the experiment in slices from mice with a homozygous GluN2B L1298A/R1300Q (GluN2B^LA/RQ^) mutation that prevents CaMKII binding^9,39^. Despite completely abolished GluN2B binding, we still saw the same level of potentiation (Figure 7B), indicating that GluN2B binding is indeed not required for the potentiation by truncated CaMKII monomers.

**Figure 7.**
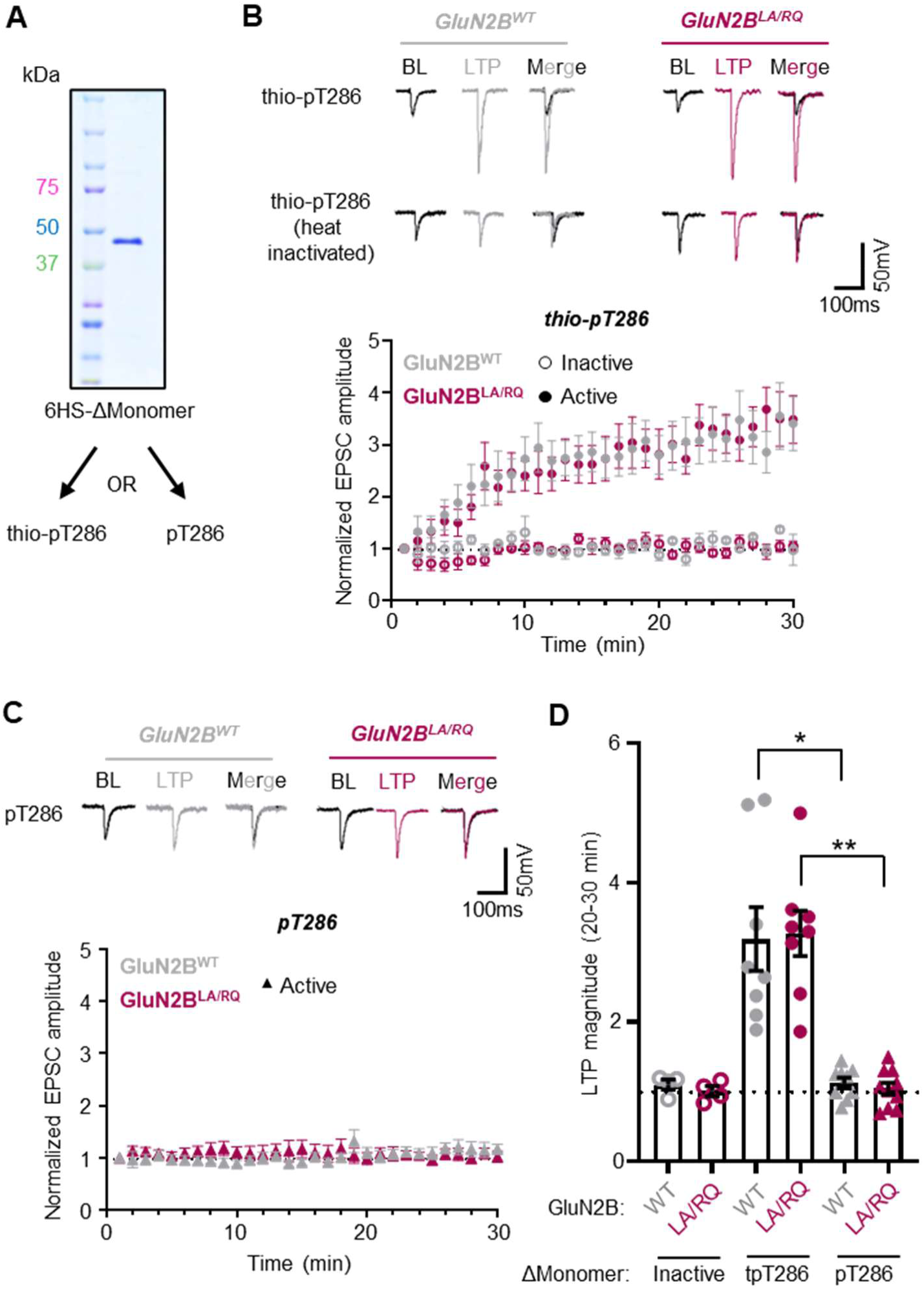
Constitutively activated CaMKII ΔMonomers cause LTP in absence of GluN2B binding. **(A)** Coomassie blue gel showing purity of purified 6xHis SUMO (6HS)-ΔMonomer (2 μM)**. (B)** Representative traces and normalized EPSC amplitude time course after infusion of 200 nM CaMKII (T286 thio-autophosphorylated -/+ heat inactivation) into CA1 of either wildtype (GluN2B WT) or GluN2B LA/RQ knock-in slices. **(C)** Representative traces and normalized EPSC amplitude time course after infusion of 200 nM CaMKII (T286 autophosphorylated) into CA1 of either wildtype (GluN2B WT) or GluN2B LA/RQ knock-in slices. **(D)** LTP magnitude measured over the final time points (20-30 min). *P<0.05; **P<0.01 by one-way ANOVA.

### A CaMKII ΔMonomer potentiates synapses only after thio-pT286 but not regular pT286

If potentiation can be induced by monomeric CaMKII even without any GluN2B binding, then why does GluN2B binding appear to be required for LTP? One possibility is that GluN2B binding is required to extend the duration of Ca^2+^-independent “autonomous” CaMKII activity. Although autonomous activity can be generated both by pT286 and by GluN2B binding^8,41^, the pT286 is sensitive to phosphatases and reversed within minutes after LTP stimuli^42–44^. Indeed, in our most recent model of LTP, the requirement for CaMKII binding to GluN2B for LTP induction^6,7^ lies in the generation of prolonged autonomous activity of the GluN2B-bound CaMKII^8^, which then mediates LTP expression^7^ (Supplemental Figure S3A). According to this model, thio-pT286 CaMKII may circumvent the requirement for GluN2B binding by its alternative mechanism to prolong autonomous activity, which cannot be as readily reversed by phosphatases as the naturally occurring regular pT286 of CaMKII. Indeed, when we repeated our experiment with the ΔMonomer using regular pT286 instead of thio-pT286 we did not detect any potentiation at all (Figure 7C). This indicates that LTP indeed requires CaMKII binding to GluN2B, but that this requirement can be overcome by extending autonomous CaMKII activity using the artificial phosphatase-resistant thio-pT286, but not by natural pT286 (Supplemental Figure S3B). The lack of potentiation by the ΔMonomer with regular pT286 was seen both in wild type slices and in slices with the GluN2B^LA/RQ^ mutation that prevents CaMKII binding (Figure 7C, D), providing further functional support for the finding that hub domain deletion is sufficient to eliminate effective GluN2B binding within cells.

## DISCUSSION

This study successfully addressed a major remaining conundrum about the CaMKII mechanisms in LTP: Why can truncated CaMKII monomers induce synaptic potentiation even though they are impaired for the GluN2B binding that is required for LTP? We here first disproved our hypothesis that this is only an apparent conundrum because GluN2B binding of truncated monomers just eluded detection: Even though GluN2B binding was indeed detected for full-length monomers that were generated by hub domain mutation, this was not the case for truncated monomers without hub domain. Further, truncated monomers still potentiated synapses even in neurons with a GluN2B mutant that abolishes CaMKII binding. Instead, we then found that GluN2B binding can be circumvented by making the monomers constitutively active in another way that is phosphatase resistant and thus prolongs the duration of kinase activity. Artificially, this can be achieved by thio-autophosphorylation (thio-pT286), T286D mutation, or deletion of the autoinhibitory region (in a 1-290 truncation)^17–19,45,46^. However, physiologically, such prolonged phosphatase-resistant autonomous activity is generated by binding to GluN2B. This supports our current overall model of LTP, in which regular CaMKII autophosphorylation (pT286) is required only during LTP induction and only to enable efficient CaMKII binding to GluN2B. Then, the autonomous activity of the GluN2B-bound CaMKII mediates LTP expression even after pT286 is reversed. Indeed, our recent studies have shown that LTP induction, expression, and maintenance can be achieved even in T286A mutant mice if the pT286 requirement for GluN2B binding is circumvented by ATP-competitive CaMKII inhibitors that directly enhance this binding^6,7^. Thus, the findings of the current study reconcile the results from several classic CaMKII studies^17–19^ with our updated modern model of the CaMKII mechanisms that mediate LTP^5–7^. Additionally, our findings emphasize that LTP expression indeed requires enzymatic activity of CaMKII, even if LTP induction may not.

Notably, although our findings support the central role of CaMKII binding to GluN2B in LTP, they suggest that the more recently described co-condensation of CaMKII with GluN2B^7,29,30^ is not required for LTP. We found that even CaMKII monomers can co-condensate with GluN2B, but this required the CaMKII hub domain that is deleted in the truncated monomers that were sufficient to induce LTP. Thus, the truncated monomers cannot form co-condensates with GluN2B, even though their thio-pT286 was sufficient to circumvent the requirement for traditional GluN2B binding. This demonstrates that synaptic potentiation can by mediated by phosphatase-resistant constitutive CaMKII activity in the absence of structural organization by co-condensation. This further indicates that physiological LTP induction and expression only require CaMKII binding to GluN2B, but not their co-condensation. As our previous studies indicated that co-condensation is not required for LTP maintenance^7^, these findings suggest that co-condensation of CaMKII with GluN2B is not required for LTP at all. While it would be beneficial to corroborate these findings with additional independent lines of evidence, this is currently not possible with any other available tools. Thus, the role of CaMKII co-condensation with GluN2B in LTP still awaits further investigation. In either case, our results do not rule out other functions of CaMKII co-condensation with GluN2B in basal synaptic organization or a function of other condensates in LTP.

The finding that full-length CaMKII monomers were able to co-condensate with GluN2B was initially surprising, as condensation requires multivalent interactions^31–33^ that were thought to require the multimeric CaMKII holoenzyme structure. However, co-condensation of full-length monomers is consistent with GluN2B interaction not only via the kinase domain (as previously shown) but also via the hub (as shown here): With multiple binding sites even on a CaMKII monomer, such monomers are sufficient to support condensation. Additionally, our studies now directly indicate multiple binding sites on GluN2B. Indeed, our co-condensation assay within cells should require multiple binding sites on GluN2B in order to work: A prior biochemical *in vitro* study used GST-GluN2Bc fusion proteins that dimerize (and thus provide multiple binding sites in the dimer even if each GluN2Bc monomer only had a single binding site). By contrast, the mScarlet-GluN2Bc fusion protein for our cellular assay should be monomeric in nature (and would thus require multiple binding sites on GluN2Bc in order to successfully engage in co-condensation).

The successful GluN2B binding of full-length CaMKII monomers may also at least partially address another remaining conundrum that we have reviewed recently^5^: CaMKII binding to GluN2B is necessary for its synaptic accumulation after LTP stimuli, yet there is much more CaMKII than GluN2B in dendritic spines. Previously, it was thought that this synaptic accumulation requires the CaMKII holoenzyme for avidity effects, i.e., for enabling simultaneous interactions of multiple CaMKII subunits of each holoenzyme with multiple GluN2B C-tails. However, with a single CaMKII subunit sufficient to mediate this binding, each bound subunit also targets the other eleven subunits of the holoenzyme to the synapse. Thus, one GluN2B subunit is sufficient to mediate the synaptic targeting of at least 12 CaMKII subunits. Additionally, the sufficiency of binding to a single GluN2B subunit explains why CaMKII still accumulates at synapses of mature hippocampal neurons, where each NMDA receptor typically contains only a single GluN2B subunit^47^. Nonetheless, additional mechanisms may be needed to explain the amount of synaptic CaMKII. Indeed, there are various other binding proteins at the synapse^4,48^, however, so far, the interaction with GluN2B C-tail is the only one shown to be both necessary for synaptic CaMKII accumulation in neurons and sufficient for reconstituting a similar movement in heterologous cells^5,38^.

The sufficiency of CaMKII monomers for both GluN2B binding and co-condensation again raises questions about the function of the multimeric CaMKII holoenzyme structure. As the results of the current study show that the CaMKII oligomerization into holoenzyme is dispensable for mediating the downstream effects of Ca^2+^-signaling (at least for LTP expression), they further bolster the notion that the CaMKII holoenzyme is instead required for the upstream processing of different Ca^2+^-signals that can lead to opposing functional outcomes in neurons, such as long-term potentiation versus depression (LTP versus LTD). Indeed, pT286 is thought to occur preferentially between neighboring subunits within holoenzymes^26–28^ and is required for induction of both LTP and LTD^49,50^. Then, the maximal pT286 that occurs in response to the strong and brief LTP stimuli promotes binding to GluN2B and accumulation at excitatory synapses^6–8,13,51,52^. By contrast, the lower level of pT286 that occurs in response to the weak but prolonged LTD stimuli instead promote additional autophosphorylation at T305/306 (pT305/306) within the holoenzyme, which suppresses further pT286 as well as the LTP-related GluN2B binding and accumulation at excitatory synapses^40,52^. Thus, CaMKII holoenzyme mechanisms are required to process LTP-versus LTD-stimuli into promotion versus suppression of GluN2B binding, even though the binding itself is sufficiently mediated by CaMKII monomers. In other words, although CaMKII monomers may be sufficient to act as downstream signaling effectors in synaptic potentiation, the CaMKII holoenzyme is required for the upstream signal computation. Although the CaMKII holoenzyme first sparked the imagination of neuroscientists as a potential device for molecular memory storage, its role in molecular signal computation may be even more intriguing.

## MATERIALS AND METHODS

### Experimental model and subject details

All animal procedures were approved by the University of Colorado Anschutz Medical Campus Institutional Animal Care and Use Committee (IACUC) and carried out in accordance with the National Institutes of Health best practices for animal use. The University of Colorado Anschutz Medical Campus is accredited by the Association for Assessment and Accreditation of Laboratory Animal Care, International (AAALAC). All animals were housed in ventilated cages on a 12 h light/12 h dark cycle and were provided *ad libitum* access to food and water. The GluN2B^LA/RQ^ mutant mouse line was kindly provided by Johannes Hell^9^. These mice and their littermate WT controls were obtained from a heterozygous breeding pair. Mixed sex pups from Sprague-Dawley rats (Charles River) were used to prepare dissociated hippocampal cultures for imaging experiments.

### Material and DNA constructs

All DNA constructs were either purchased from AddGene or received as a gift and were further cloned to include the indicated fluorophores, mutations, and truncations. Gibson assembly was used for changes to fluorophores and N-terminal truncations. Site-directed mutagenesis by PCR was used to generate all point mutants and C-terminal truncations. All plasmids were fully sequenced prior to their use in experiments.

### HEK293T cell GluN2B binding assay

HEK293T cells were plated in a 12-well plate on #1 coverslips and transfected by the Ca^2+^ phosphate method at ∼50% confluency with 1.5 μg of each of the following plasmids (unless specified otherwise): mEGFP-CaMKII (and mutants), pDisplay-mCherry-GluN2B, and pNES-iRFP-C1 as a cell fill. 24 hours following transfection, cells were imaged at 34°C in a HEPES-buffered imaging solution containing (in mM): 130 NaCl, 5 KCl, 10 HEPES (pH 7.4), 20 glucose, 2 CaCl2, and 1 MgCl2. Live, healthy cells with expression of all desired plasmids were selected for imaging. All images were collected with a spinning disc confocal microscope with a 63X oil immersion objective. Baseline images were collected, then ionomycin (10 μM) was used to induce Ca^2+^ influx and activate CaMKII. Images were then taken every minute until 5 minutes post-ionomycin treatment. Cells were analyzed in FIJI ImageJ for mEGFP/mCherry co-localization by Pearson’s correlation within a cytoplasm mask based on the cell fill. Each data point represents quantification of a single cell.

### Protein expression and purification

Fluorescently tagged CaMKII for fluorescence correlation spectroscopy and *in vitro* GluN2B binding was expressed for 48 hours in HEK293T cells. Media was removed and replaced with ice cold phosphate buffered saline and cells were scraped with a rubber policeman and pelleted at 2500 RPM at 4°C for 5 min. The supernatant was discarded, and pellets were resuspended in homogenate buffer containing (in mM) 50 PIPES pH 7.12, 1 EGTA, 1 DTT, 10% glycerol and complete protease inhibitor cocktail (Roche). The homogenate was then spun at 14,000 RPM for 20 min at 4°C to remove cell debris and the resulting supernatant was collected. A western blot was used to quantify [CaMKII] by using a standard curve made with purified CaMKII. Lysates were diluted with mock-transfected cell lysate to achieve equal [CaMKII] between samples.

GST-GluN2B and mutants were expressed and purified from *E. Coli* BL21 DE3 cells as previously described^15,20^. Briefly, protein expression was induced for 2 hours at room temperature with 1 mM IPTG. Cells were resuspended in 20 mM Tris pH 7.55, 150 mM NaCl, 0.1% Tween-20, 1 mg/mL lysozyme, 1X protease inhibitor cocktail, MgCl_2_, DNase, and RNase on ice for 15 minutes. The resuspended cells were sonicated and ultracentrifuged at 4°C for 30 min at 10,000xg. The resulting pellet was resuspended in a buffer containing STE (20 mM Tris, 0.15 M NaCl, 1 mM EDTA) and 1X protease inhibitor cocktail. DTT (1 mM) and sarcosyl (10%) were added to the resuspensions which were then sonicated and ultracentrifuged at 4°C for 20 min at 10,000xg. 3% Triton X-100 was added to the supernatant, which was then added to Glutathione Sepharose (GE Healthcare) for batch purification and eluted with 100 mM reduced glutathione in 200 mM Tris, pH 9.0. The protein was dialyzed for 3 hours at room temperature in 200 mM Tris, pH 9.0 to remove glutathione.

CaMKII constructs were expressed with N-terminal 6xHis-SUMO (6HS) tags in Rosetta2 cells based on a published protocol^16^. Briefly, constructs were co-expressed with λ-phosphatase and induced with IPTG at 18°C overnight. Cells were lysed for 20 min on ice in a buffer containing: 25 mM Tris-HCl, 150 mM KCl, 50 mM imidazole, 10% glycerol, 25 mM MgCl_2_, 1X EDTA-free protease inhibitor cocktail, 1 ug/mL DNAse, 0.2 mM AEBSF, 1 mg/mL lysozyme, 1:1000 Tween20, pH 8.5. After three rounds of 10 seconds of sonication at 50% power, the lysate was ultracentrifuged at 100,000xg for 30 minutes at 4°C. The soluble lysate loaded onto a Nickel Sepharose column (Cytiva) using a Knauer Azura FPLC system and eluted with a buffer containing 25 mM Tris-HCl, 150 mM KCl, 500 mM imidazole, 10% glycerol, 500 mM imidazole pH 8.5. Purity of the eluted samples was confirmed by SDS-PAGE followed by Coomassie staining.

### Fluorescence correlation spectroscopy

Fluorescence correlation spectroscopy (FCS) analysis was performed on a Zeiss LSM780 spectral microscope equipped with ZEN2011 software and an FCS/RICS package. 50 nM fluorescein was used to optimize the microscope pinhole position and to determine the structural parameter of the detection volume. Fluorescent protein tagged CaMKII extracts from HEK293T cells were diluted to achieve a baseline count rate of 1– 100 kHz to achieve an average particle number (N) of ≤10 in the focal volume. Ten acquisitions were performed for 10 seconds each and averaged together for a single n, represented by a single point on each graph. Traces with obvious indications of protein aggregates were excluded from molecular brightness analysis. Autocorrelation curves were fitted with a 3D Gaussian distribution model using PyCorrFit software. Molecular brightness is reported as counts per particle (kHz). Diffusion time is reported as residence time in focal volume (ms).

### HEK293T cell light-induced co-clustering (LINC) assay

HEK293T cells were plated and transfected as done in the HEK293T cell GluN2B binding assay. Cells expressing mEGFP-CaMKII, CIBN-mCherry-CaMKII, and CRY2olig were first imaged in the dark to exclude any cells with pre-light clustering. 488 nm light was used to induce clustering, and additional post-light images were acquired at least one minute later. Clusters were identified by intensity of at least two standard deviations above the average fluorescence of the cell. Masks were then made for the clustered and non-clustered mCherry regions of the cell. Quantifications are expressed as mEGFP clustered intensity / mEGFP non-clustered intensity (“mEGFP clustered/non-clustered”). Each data point represents quantification of clusters in a single cell.

### In vitro kinase reactions

HEK cell lysates were diluted with lysates from mock-transfected cells to achieve the desired concentration for each of the mEGFP-CaMKII constructs, which was estimated by western blot. Kinase reactions were performed with 5 nM total CaMKII subunit concentration on ice (0°C) for 1 minute in the presence of: 50 mM PIPES, 2 mM CaCl_2_, 10 mM MgCl_2_, 1 mM ATP, 1 μM calmodulin, and 2 μM microcystin. Reactions were stopped by addition of a sample buffer containing 54 mM Tris pH 6.8, 1.6% SDS, 8% glycerol, 50 mM dithiothreitol, and 0.013% bromophenol blue. Western blot was used to measure total CaMKII (with a homemade CBα2 antibody) and pT286 (Phosphosolutions).

### Western blot

Protein was separated by SDS-PAGE and transferred onto low-fluorescence PVDF membranes for 1 hour at 4°C. Membranes were blocked at room temperature for 30 min with a 50/50 (vol/vol) mixture of TBS-T and Fluorescent Blot Blocking Buffer (Azure) and incubated overnight at 4°C in primary antibodies in the same blocking buffer. Protein was detected by primary antibodies against: CaMKIIα (1:1000), pT286 (1:2000; Phosphosolutions) GST (1:1000), or GFP (1:1000). An anti-GFP antibody against the first 18 amino acids (that are identical between similar fluorophores) was used for equal detection of protein tagged with different fluorophores. Blots were next incubated with fluorescent secondary antibodies (Azure) diluted 1:10,000 in blocking buffer at room temperature for 45 min. Blots were developed by fluorescence detection with an Azure imager and analyzed for densitometry in FIJI ImageJ. Molecular weight markers are color coded based on the Biorad Kaleidoscope Standard.

### In vitro GluN2B binding

CaMKII-GluN2B binding was tested with GST-GluN2B constructs as indicated in figures. Each lot of anti-GST plates were first tested to determine the amount of GST-GluN2B required to saturate the anti-GST antibody. After this initial characterization, the appropriate amount of GST-GluN2B (diluted in PST buffer containing 0.05 % BSA) was bound to the microtiter plate for one hour with gentle agitation at room temperature and washed three times with PST. The plate was then blocked in 5% BSA in PST for 30 minutes with gentle agitation at room temperature and washed one time with PST. HEK293T extracts containing CaMKII at the indicated concentrations were bound for 30 min in CaMKII binding buffer containing 50 mM PIPES pH 7.1, 150 mM NaCl, 1 mM MgCl_2_, 100 μM ADP, 0.05% BSA, 0.05% Tween-20, and Ca^2+^/CaM (2 mM/1 μM). To assay Ca^2+^-independent binding, the Ca^2+^/CaM was replaced with 1 mM EGTA. Unbound CaMKII was washed away in PST containing 1 mM EGTA 4 times for 3 minutes each. Western blot loading buffer containing 1.6% SDS was then added to the wells and incubated at 95°C for 10 minutes. Eluted samples were then analyzed by SDS-PAGE and western blot. Each data point represents an individual well of the microtiter plate.

### Primary hippocampal neuronal cultures

Mixed sex Sprague Dawley rat pups were used for hippocampal dissection on postnatal day 0 (P0). Dissected hippocampi were dissociated in papain solution for 1 hour and then plated at a density of 100,000 cells/mL on 18 mm #1 coverslips in plating media (MEM containing 10% FBS, 1% Penicillin-Streptomycin) in 12-well plates. On day *in vitro* (DIV) 1, the plating media was replaced with feeding media (Neurobasal A containing 2% B27 and 1% Glutamax). On DIV 7, half of the conditioned media was replaced with fresh feeding media containing 2% 5-Fluoro-2’-deoxyuridine to limit growth of glial cells. Neuron cultures were maintained at 37°C with 5% CO_2_.

### Live imaging of CaMKII in neurons

DIV14-16 dissociated hippocampal neurons were transfected by the Lipofectamine method with 1 μg of each the following plasmids per well of a 12-well plate (unless specified otherwise): shRNA-CaMKIIα, mEGFP-CaMKII (and mutants), PSD95-intrabody, and HaloFill. To visualize HaloFill, 1 uL of JaneliaFluor635 was added to the neuron culture 5 minutes prior to imaging and was washed out upon adding the neuron coverslip to the imaging chamber in imaging solution. Live imaging of cultured rat hippocampal neurons occurred in a climate-controlled chamber at 34°C in a HEPES buffered artificial cerebrospinal fluid (aCSF) solution containing the following (in mM): 130 NaCl, 5 KCl, 10 HEPES pH 7.4, 20 glucose, 2 CaCl_2_, and 1 MgCl_2_. Chemical LTP (cLTP) was induced using 100 μM glutamate and 10 μM glycine. After 30 sec of cLTP treatment, 4 volumes of fresh aCSF were slowly perfused through the imaging chamber to wash out the stimulation for a total of 30 sec. Images were quantified before and immediately after washout of cLTP every minute for 5 minutes. Spines were identified by expression of an intrabody against a marker protein for excitatory synapses, PSD-95. A mask was made for the “synapse” (i.e., PSD95-positive region) and for the “shaft” (i.e., HaloFill minus synapse). The intensity of CaMKII is quantified in both masks and expressed as the synapse/shaft ratio (“Synaptic CaMKII”). Each individual point represents the average synaptic CaMKII value for one ROI (one stretch of dendrite).

### HEK293T cell GluN2B co-condensation assay

HEK293T cells were plated onto glass coverslips pre-coated with 0.1 mg/mL poly-D-lysine (PDL) and transfected using the calcium phosphate method as previously described^7,14^. In this assay, mEGFP-CaMKII was co-expressed with a soluble, non-membrane-targeted mScarlet-tagged GluN2B C-terminal tail (mScarlet-GluN2Bc). 24-30 hours post-transfection, live-cell imaging was conducted at 34 °C in a HEPES-buffered imaging solution containing: 130 mM NaCl, 5 mM KCl, 10 mM HEPES (pH 7.4), 20 mM glucose, 2 mM CaCl_2_, and 1 mM MgCl_2_. Co-condensate formation was triggered by a calcium stimulus induced by bath-application of 10 μM ionomycin for 6 minutes. To assess the stability of condensates, 2.5 mM EGTA was subsequently bath applied for 5 minutes to chelate intracellular Ca²⁺. Quantification of condensate dynamics was performed as previously described^7,35^. The CaMKII intensity ratio was defined as the fluorescence intensity of mEGFP-CaMKII within puncta divided by the total mEGFP-CaMKII fluorescence in the cell at each time point. Cells with either very low or very high CaMKII expression were excluded to ensure that comparisons were made between cells with comparable, intermediate expression levels.

### In vitro thio-autophosphorylation

Purified 6HS-CaMKII constructs were made constitutively active by *in vitro* autophosphorylation, similar as previously described in one of the classic studies that suggested sufficiency of enzymatic CaMKII activity for synaptic potentiation^18^. Briefly, a 2 μM stock of purified protein was incubated for 10 minutes on ice in the presence of 50 mM HEPES, 2 mM CaCl_2_, 10 mM MgAcetate, 1 mM ATP OR ATPγS (for autophosphorylation OR thio-autophosphorylation), 6 µM calmodulin, and 1 mg/mL BSA. Heat inactivated protein was boiled at 100°C following thio-autophosphorylation. All forms of purified protein were diluted 1:10 to achieve a final concentration of 200 nM in the internal solution for electrophysiology experiments. The Ca^2+^-independent autonomous activity of autophosphorylated and thio-autophosphorylated kinase was confirmed by *in vitro* kinase activity assay.

### Preparation of hippocampal slices

Male mice (3-5 weeks) were anesthetized with isoflurane, decapitated, and their brains were rapidly dissected. 300 μm horizontal slices containing the hippocampus were sectioned with a vibratome (Leica VT1200) in ice cold high-sucrose cutting solution containing (in mM) 85 NaCl, 75 sucrose, 25 D-glucose, 24 NaHCO_3_, 4 MgCl_2_, 2.5 KCl, 1.3 NaH_2_PO_4_, and 0.5 CaCl_2_. Slices were transferred to 31.5°C oxygenated aCSF containing (in mM) 126 NaCl, 26.2 NaHCO_3_, 11 D-Glucose, 2.5 KCl, 2.5 CaCl_2_, 1.3 MgSO_4_-7H2O, and 1 NaH_2_PO_4_ for 30 min, then recovered at room temperature for at least 1 hour before recordings.

### Electrophysiology recordings

Whole-cell patch-clamp recordings were made at 29.5°C aCSF, with 3-5 MΩ patch pipettes, and cells were voltage-clamped at -70mV. All recordings were acquired using Molecular Devices Multiclamp 700B amplifier and Digidata 1440 digitizer with Axon pClamp™ 9.0 Clampex software, lowpass filtered at 2 kHz, and digitized at 10 kHz. A potassium gluconate internal solution containing (in mM) 137 K-gluconate, 5 KCl, 10 HEPES, 10 Phosphocreatine, 4 Mg_2_-ATP, 0.5 Na_2_-GTP, and 0.2 EGTA with purified CaMKII (as described above) was used for measuring evoked excitatory postsynaptic currents (EPSCs). EPSCs were isolated pharmacologically with 100μM picrotoxin in the aCSF. Synaptic responses were evoked by stimulation with a homemade nichrome electrode placed in the stratum radiatum (A-M Systems 2100 Isolated Pulse Stimulator). CA3-CA1 EPSCs were evoked at 0.1 Hz for 30 min. Notably, the original study^18^ included the phosphatase inhibitor microcystin in the patch pipette to obtain consistent results, despite using thio-pT286. By contrast, our measurement did not include any phosphatase inhibitor, neither for the thio-pT286 nor the regular pT286 CaMKII. In fact, in our initial experiment that did include phosphatase inhibitors (either microcystin or okadaic acid), we were unable to keep cells alive long enough to obtain any stable recordings at all.

### Analysis of electrophysiology recordings

EPSC peak amplitudes from each recording were identified using Axon™ pClamp10 Clampfit software. To represent the data, 6 sweeps (0.1 Hz) were averaged at each minute and normalized to the first minute of recording. The magnitude of the overall synaptic potentiation was expressed as the average amplitude of EPSCs for the final 10 min of the recordings. Cells of poor quality were excluded from electrophysiology analysis (i.e., access resistance > 20% of the membrane resistance or changed >10% during experimentation).

### STATISTICAL ANALYSIS

Data are shown as mean ± SEM unless indicated otherwise. All data were tested for ability to meet parametric assumptions including normality, homoscedasticity, and independence. Statistical significance is indicated in the figure legends and n’s are shown as individual data points on graphs. Individual methods sections for each technique indicate the meaning of each individual data point. Data were analyzed using Prism (GraphPad) software. All data were tested for parametric conditions, as evaluated by a Shapiro-Wilk test for normal distribution and a Brown-Forsythe test (3 or more groups) or an F-test (2 groups) to determine equal variance. Comparisons between two groups were analyzed using unpaired, two-tailed Student’s t tests. Comparisons between pre-and post-treatment images from the same cells were analyzed using paired, two-tailed Student’s t tests. Comparisons between more than two paired treatment conditions were made by repeated measures (RM) ANOVA. Comparisons between three or more groups were made by one-way ANOVA with post-test by Bonferroni, unless otherwise indicated. Statistical significance is indicated, including by *P<0.05; **P<0.01; ***P<0.001, or ns, not significant.

## ACKNOWLEDGEMENTS

We thank Dr. Meg Stratton (UMass Amherst) for kindly sharing the 6HS-CaMKIIα construct. We also thank Dr. Petrine Wellendorph and Dr. Bente Frølund (University of Copenhagen) for kindly providing the HOCPCA compound. This work was supported by National Institutes of Health grants T32 GM007635 (supporting C.N.B., C.M.B., and C.N.M.), F31 NS129254 (to C.N.B.), T32 GM008497 (supporting C.N.M.), F30 DA057053 (to C.N.M.), R21 MH140328 (to J.A.), R01 NS081248, R01 NS118786, and R01 AG067713 (to K.U.B.).

## AUTHOR CONTRIBUTIONS

C.N.B., C.M.B, C.N.M., and S.J.C. and performed experiments and analyses; C.N.B. and K.U.B. conceived this study. C.N.B. and K.U.B. wrote the original draft of this paper and all authors contributed to the final version of the manuscript.

## DECLARATION OF INTERESTS

K.U.B. is co-founder and board member of Neurexis Therapeutics, a company that seeks to develop a CaMKII inhibitor into a therapeutic drug for cerebral ischemia. C.N.B., S.J.C., and K.U.B. are named inventors on patent applications related to CaMKII inhibitors that were submitted by the Regents of the University of Colorado.

**KEY RESOURCES**

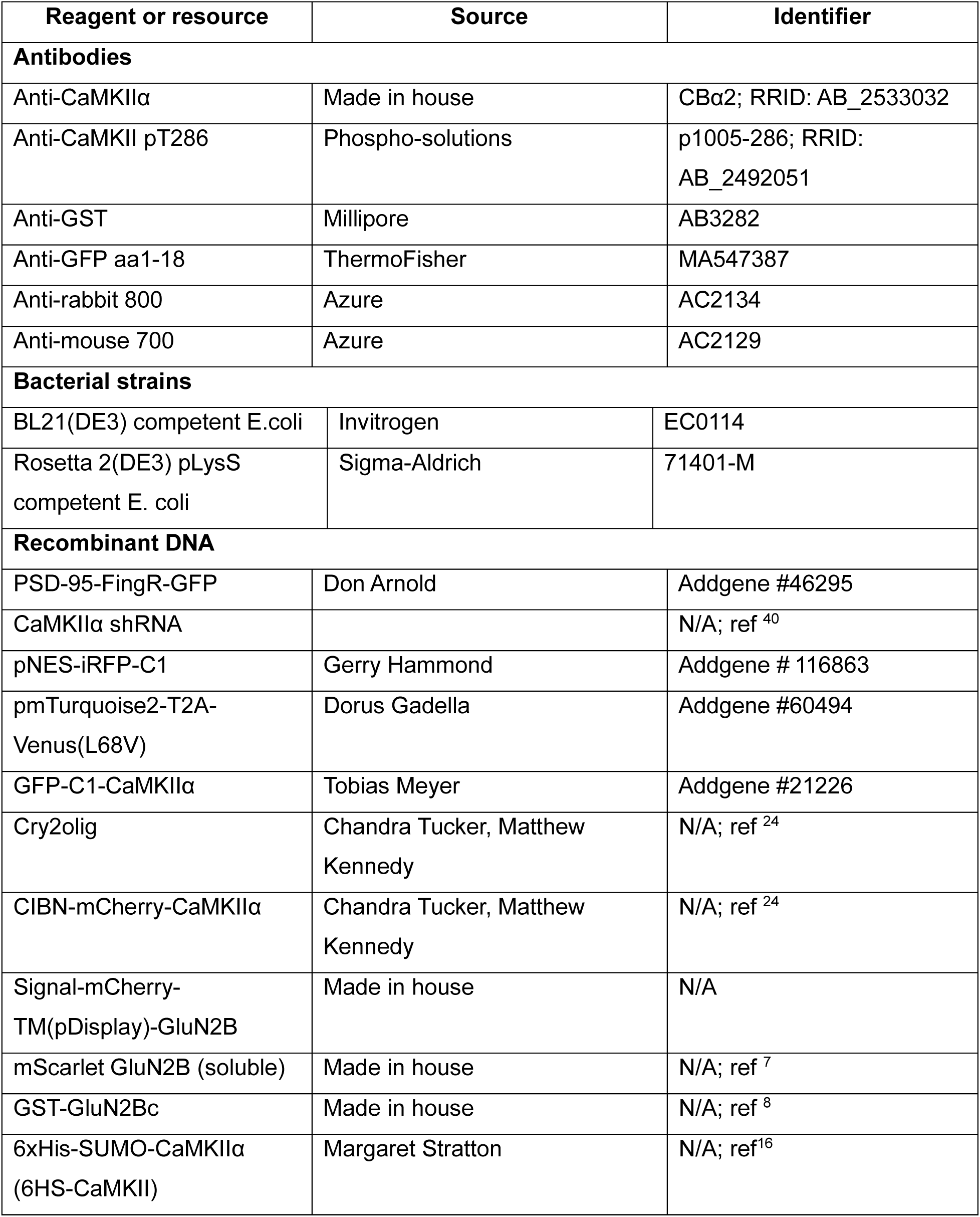
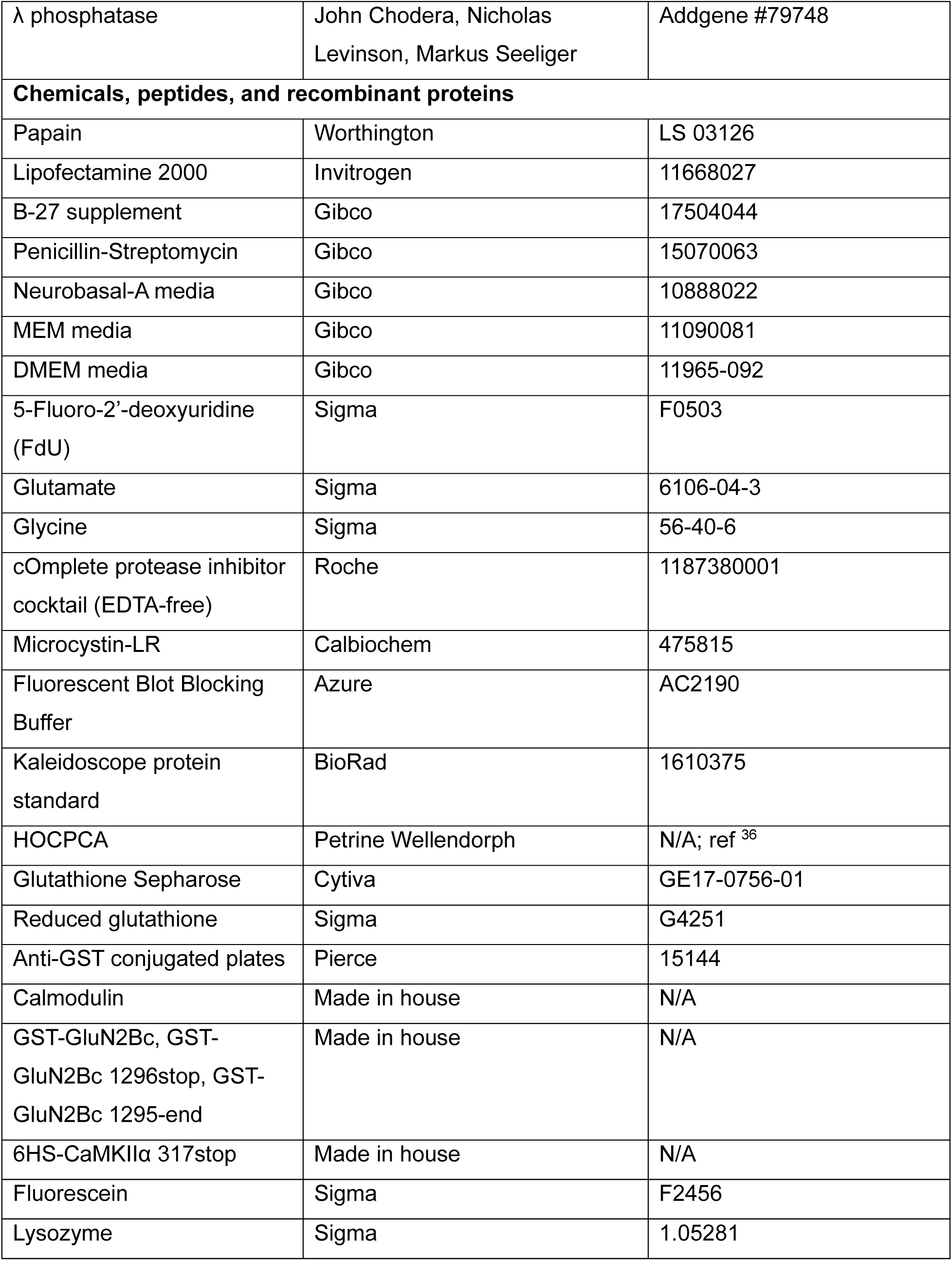
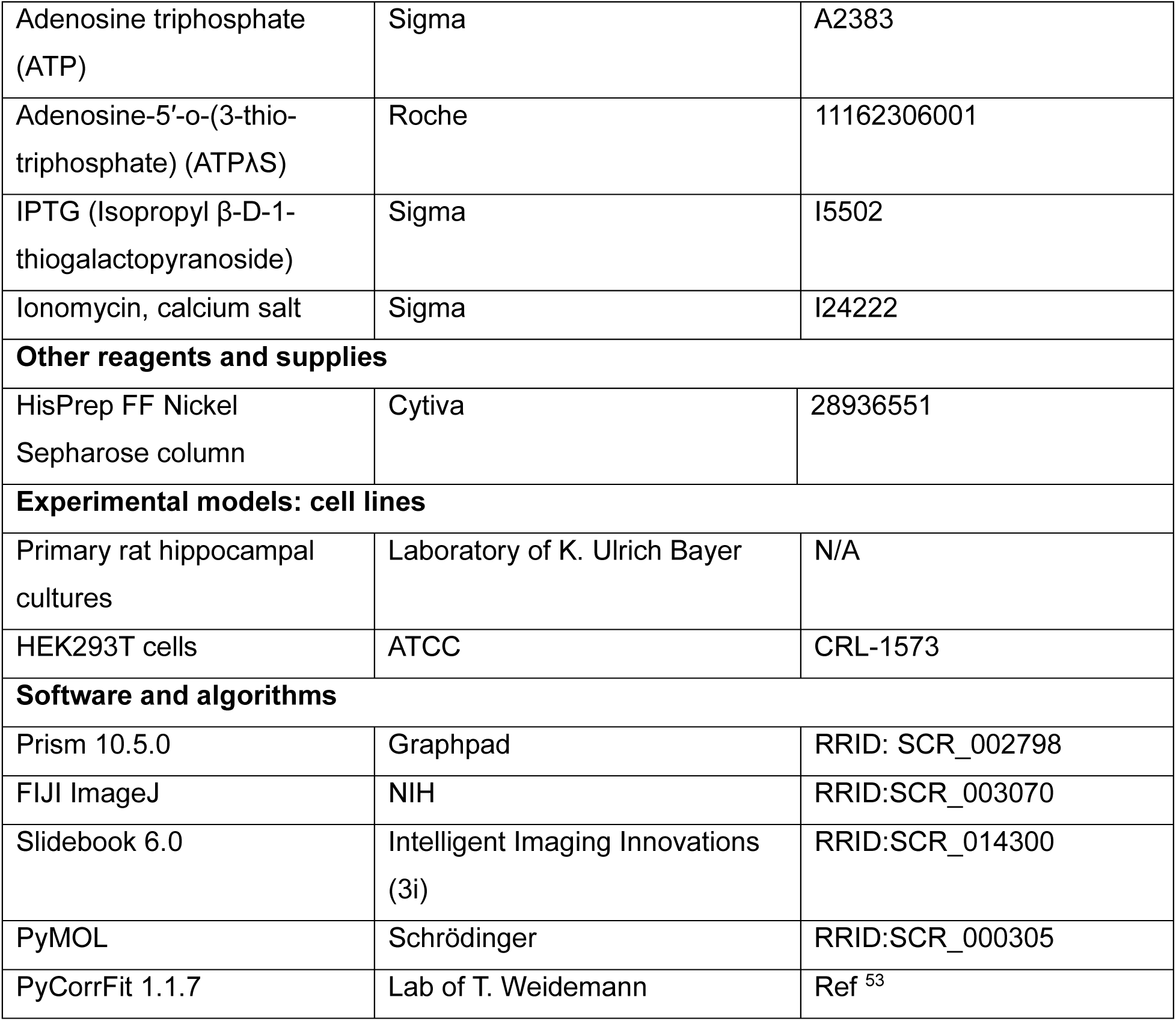

