## Supplemental Figures for "CaMKII monomers are sufficient for GluN2B binding, co-condensation, and synaptic potentiation"

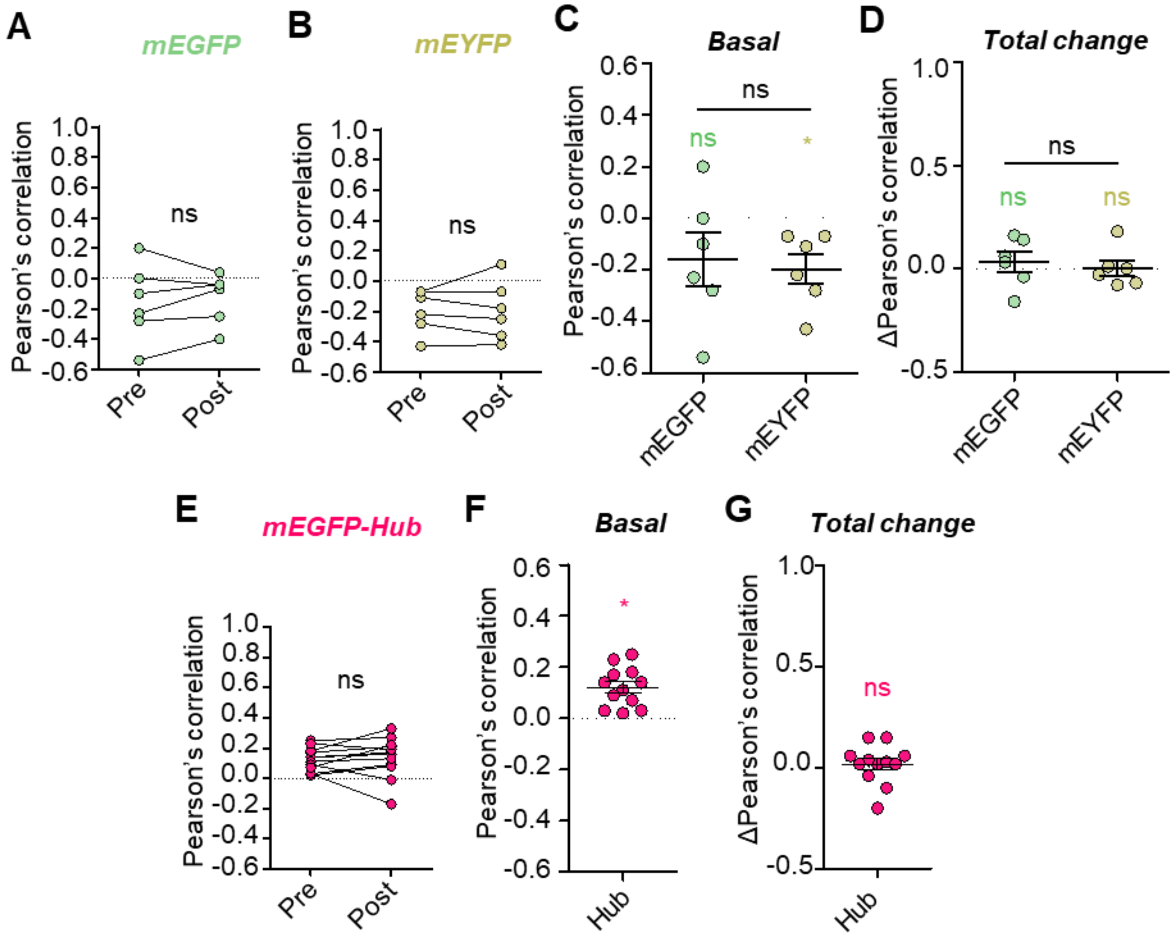


**Supplemental Figure S1. No basal or ionomycin-induced co-localization of fluorescent protein tags with GluN2B.** Paired quantification of **(A)** mEGFP or **(B)** mEYFP co-localization with pDisp-mCh-GluN2B pre and 5 minutes post ionomycin stimulation in HEK cells. ns, not significant by paired t-test. **(C)** Basal co-localization of mEGFP or mEYFP with pDisp-mCh-GluN2B. Colored symbols: *$P<0.05$; ns, not significant vs 0 by one sample t-test. Black symbols: ns, not significant by unpaired t-test. **(D)** Total change (post – pre) of pDisp-mCh-GluN2B co-localization. Colored symbols: ns, not significant versus 0 by one sample t-test. Black symbols: ns, not significant by unpaired t-test. **(E)** Paired quantification of mEGFP-CaMKII hub with pDisp-mCh-GluN2B pre and 5 min post ionomycin stimulation. ns, not significant by paired t-test. **(F)** Basal co-localization of mEGFP-CaMKII hub with pDisp-mCh-GluN2B. Colored symbols: *$P<0.05$ vs 0 by one sample t-test. **(G)** Total change (post – pre) of pDisp-mCh-GluN2B co-localization. ns, not significant versus 0 by one sample t-test.

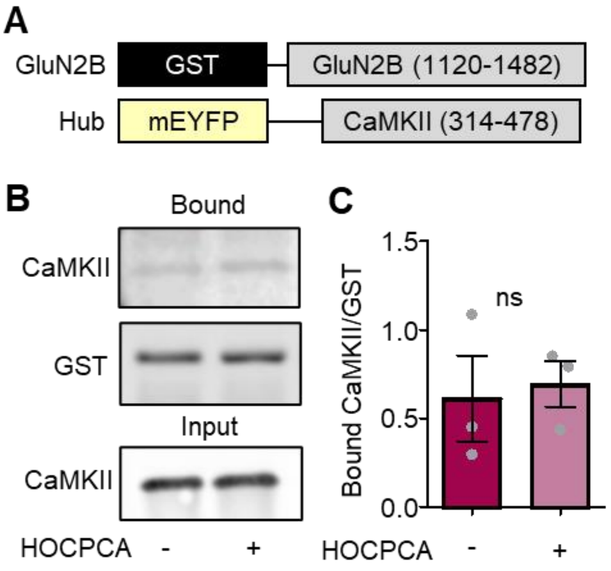


**Supplemental Figure S2. No effect of HOCPCA on CaMKII hub GluN2B binding *in vitro*.** **(A)** Constructs used in this experiment. **(B)** Representative blots and **(C)** quantification. Binding of the CaMKII hub to GluN2B in the presence of $Ca^{2+}$/CaM (2 mM/1 μM) and ADP (100 μM) is not affected by HOCPCA (2 mM). ns, not significant by unpaired t-test.

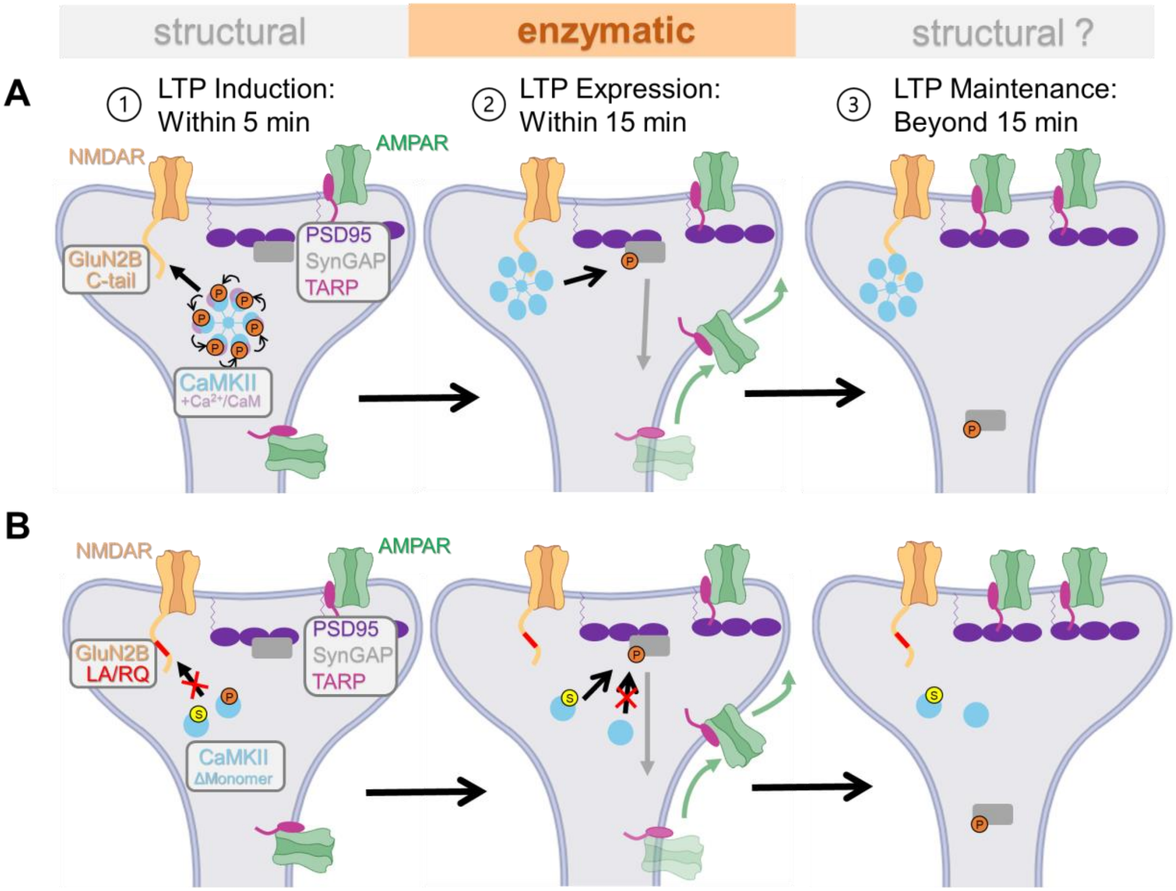


**Supplemental Figure S3**. **Model of CaMKII functions in LTP. (A)** (1) Induction of physiological LTP requires CaMKII binding to GluN2B, which is positively regulated by autophosphorylation at T286 (orange P). This GluN2B binding then generates the $Ca^{2+}$-independent autonomous activity of CaMKII required for (2) LTP expression, even upon T286 dephosphorylation. **(B)** The requirement for such GluN2B-derived autonomous activity during (2) LTP expression can be circumvented by artificially prolonging CaMKII activity, for example, by T286 thio-autophosphorylation (yellow S).
